# Human RAMP1 overexpressing mice are resistant to migraine therapies for motion sensitivity

**DOI:** 10.1101/2023.10.24.563838

**Authors:** Shafaqat M. Rahman, Linda Guo, Carissa Minarovich, Laura Moon, Anna Guo, Anne E. Luebke

## Abstract

Both enhanced motion-induced nausea and increased static imbalance are observed symptoms in migraine and especially vestibular migraine (VM). Motion-induced nausea and static imbalance were investigated in a mouse model, nestin/hRAMP1, expressing elevated levels of human RAMP1 which enhances CGRP signaling in the nervous system, and compared to non-affected littermate controls. Behavioral surrogates such as the motion-induced thermoregulation and postural sway center of pressure (CoP) assays were used to assess motion sensitivity.

Nausea readouts revealed that the nestin/hRAMP1 mouse exhibit an increased sensitivity to CGRP’s effects at lower doses compared to unaffected controls. In addition, the nestin/hRAMP1 mice exhibit a higher dynamic range in postural sway than their wildtype counterparts, along with increased sway observed in nestin/hRAMP1 male mice that was not present in male unaffected controls. Results from migraine blocker experiments were challenging to interpret, but the data suggests that olcegepant is incapable of reversing CGRP-induced or endogenous alterations in the nestin/hRAMP1 mice, while rizatriptan was ineffective in both the nestin/hRAMP1 and control mice. The results indicate that overexpression of hRAMP1 leads to heightened endogenous CGRP signaling. Results also suggest that both olcegepant and rizatriptan are ineffective in reducing nausea and sway in this hypersensitive CGRP mouse model. This study suggests that the hypersensitive nestin/hRAMP1 mouse may serve as a model for difficult to treat cases of migraine that exhibit increased motion sensitivity.

## Introduction

Migraine is a chronic neurological disorder that is marked by headache pain, nausea, and heightened sensitivity to environmental stimuli such as light and sound. Certain people may also experience an aura before the onset of migraine symptoms ^1^. Approximately 50% of patients with a history of migraine may experience recurrent vertigo and imbalance ^2–6^.

Both migraine and migraine with vertigo or vestibular migraine (VM) affects about 4% of adults in the US, with three times as many females affected as males. Furthermore, individuals with VM and migraine exhibit greater postural sway when encountering challenging surroundings and are more prone to motion-induced nausea then healthy subjects. ^7–12^.

Calcitonin gene-related peptide (CGRP) is commonly considered a crucial factor that causes migraines. CGRP is a vasoactive peptide present in the nervous system that contributes to vasodilation, inflammation, and pain modulation. CGRP signals through the CGRP receptor consisting of three subunits: calcitonin-like receptor (CLR), receptor activity-modifying protein 1 (RAMP1), and receptor component protein (RCP). The RAMP1subunit is essential for ligand binding and CLR translocation to the membrane and may also be the rate-limiting subunit ^13–15^.

A transgenic mouse model (nestin/hRAMP1)created by Russo et al. manifests sensitivity to CGRP by upregulating the CGRP receptor ^14^. This nestin/hRAMP1 mouse model shows increased hRAMP1 expression in the nervous system and heightened sensitivity to the neuropeptide CGRP, as compared to wildtype mice. The mice were subjects of preclinical studies that analyzed surrogate behaviors for photophobia and allodynia. Their aversive reaction to light is amplified by intracerebroventricular (ICV) CGRP injections ^14, 16, 17^. Moreover, hind paw withdrawals were detected at markedly lower doses of CGRP compared to their respective controls ^18^.

The CGRP receptor antagonist olcegepant (BIBN4096) effectively alleviates CGRP- induced behaviors in both human and mouse receptors. Moreover, preclinical models show that light aversion caused by CGRP is reduced through selective serotonin receptor agonists like rizatriptan and sumatriptan. Early studies targeting CGRP with olcegepant have led to the development of improved "gepants" and monoclonal antibodies that can block either the CGRP receptor or the peptide. Numerous treatment options for migraines based on antagonizing CGRP signaling have been investigated in clinical trials, many of which have been approved by the FDA since 2018, and a few studies have indicated that CGRP antagonists have been successful in treating migraine with vertigo ^19, 20^. Alternative treatments, such as vestibular suppressants, have not been shown to alleviate vertigo and may cause drowsiness ^21–23^. Therefore, further research is needed to investigate the potential role of CGRP sensitization in causing vestibular symptoms and to elucidate the associated mechanisms. CGRP is widely distributed throughout the vestibular nervous system (CNS and PNS)) and is present on vestibular efferent fibers that innervate the inner ear ^24–26^. Because individuals with migraine exhibit greater postural sway and are more prone to motion-sickness ^7–12^, we decided to assess these behaviors in the nestin/hRAMP1 mouse model. These behaviors were chosen as they exhibit excellent test-retest reliability, require no training, and have excellent translational capabilities ^27, 28^.

Therefore, in this study, we aim to characterize the effects of CGRP in nestin/hRAMP1 mice by employing postural sway assays, and motion-induced thermoregulation-a nausea surrogate assay. Furthermore, we seek to determine whether olcegepant or rizatriptan can provide protection against any CGRP-induced or endogenous changes in the hypersensitized CGRP receptor mouse.

## Materials & Methods

### Animals

A total of 175 mice (95F/80M) were tested across all experiments, and all studies were sufficiently powered to detect sex differences. Mice were housed under a 12 to 12 day/night cycle at the University of Rochester’s Vivarium under the care of the University of Rochester’s Veterinary Services personnel. All animal procedures were approved by the University of Rochester’s IACUC committee and performed in accordance with NIH standards.

The nestin/hRAMP1 mice were bred by crossing heterozygous or homozygous hRAMP1 loxP mice with heterozygous nestin-Cre (JAX # 3771) mice to produce offspring that express elevated hRAMP1 in the nervous system^15^. Unaffected littermate control mice used in this study were hRAMP1 loxP mice not expressing nestin-Cre. Mice were housed under a 12-hour day/night cycle under the care of the University Committee on Animal Resources (UCAR) at the University of Rochester. Mice were housed with ad libitum access to necessities (food, water, bedding, and enrichment) . Mice were equilibrated in a testing room with an ambient temperature between 22-23°C for at least 30 minutes prior to testing. Mice were tested between 3-6 months of age. Different cohorts of mice were used to test motion-induced nausea and postural sway. Testing occurred from 9:00 am to 5:30 pm.

### Drug administration

All intraperitoneal (IP) injections were performed with a fine 33-gauge insulin syringe. Dulbecco PBS (saline) served as vehicle control diluent. The CGRP-receptor antagonist olcegepant and the selective serotonin receptor agonist rizatriptan were used as migraine blockers in this study. The concentrations are listed: 0.1x, 0.5x, and 1.0x CGRP were prepared at 0.01, 0.05, and 0.1 mg/kg (rat ɑ-CGRP, Sigma), 1.0x olcegepant was prepared at 1.0 mg/kg (BIBN4096, Tocris), and 1.0x rizatriptan was prepared at 0.6 mg/kg (Sigma-Aldrich). Mice were tested approximately twenty to thirty minutes after IP injections. Animals were gently handled without the need for anesthesia. All animal procedures were approved by the University of Rochester’s (IACUC) and performed in accordance with the standards set by the NIH.

**Fig. 1** provides details regarding experimental methodologies pursued in this study.

**Fig. 1:**
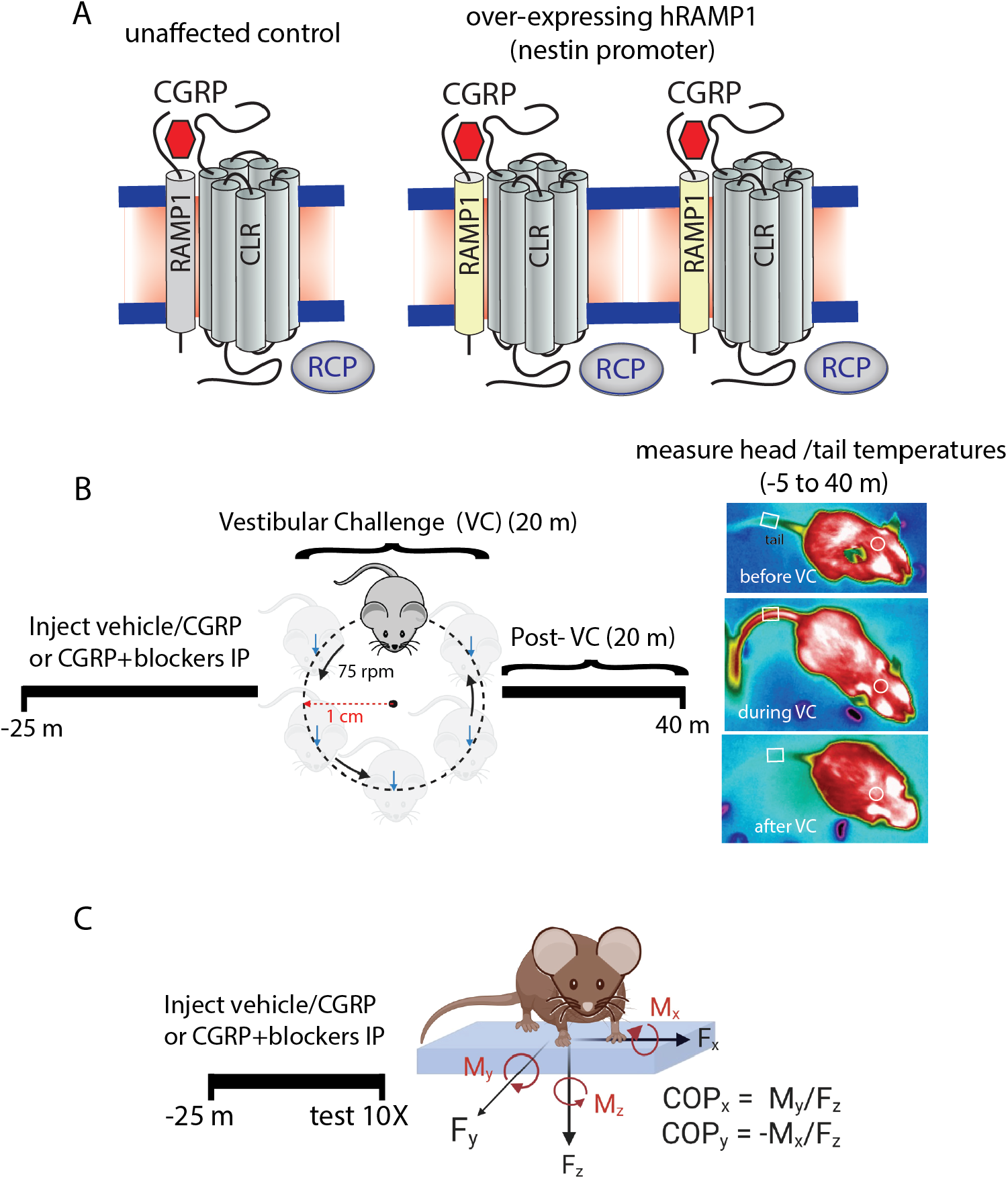
(A) Calcitonin gene-related peptide (CGRP) receptor structure consists of receptor activity modifying protein 1 (RAMP1), calcitonin-receptor like receptor (CLR), and receptor component protein (RCP). The nestin/hRAMP1 mice are designed to have enhanced CGRP signaling due to overexpression of human RAMP1 (hRAMP1). **(B-C)** Methods for motion- induced thermoregulation and center of pressure (CoP) testing as surrogate behaviors for motion-induced nausea and postural sway.

### Motion-induced nausea testing

We used a FLIR E60 IR camera (model: E64501) to head and tail temperatures of mice before, during, and after a vestibular perturbation. The measurements were conducted over a duration of 45 minutes. To summarize, we recorded baseline measurements for five minutes prior to initiating the perturbation (-5 mins ≤ t ≤ 0 mins). Subsequently, mice were recorded for 20 minutes (0 mins ≤ t ≤ 20 mins) during an orbital rotation at 75 rpm (2 cm orbital displacement). After the perturbation, mice were additionally recorded for 20 minutes to observe the recovery from hypothermia back to baseline (20 min ≤ t ≤ 40 min). FLIR Tools+ was used for analyzing data. We defined Δ tail vasodilations (°C) as the transient increases in mouse tail temperature in response to perturbation. These were computed by subtracting the baseline tail temperature at time t = 0 minutes from the maximum tail temperature measured during the first 10 minutes of the rotation (0 ≤ t ≤ 10). Additionally, we calculated the hypothermia magnitude (Δ heads) by subtracting the baseline head temperature at time t = 0 minutes from the minimum head temperature recorded throughout the entire experiment (0 mins ≤ t ≤ 40 mins). This nausea surrogate assay exhibits excellent test-retest reliability and only shows temperature changes with provocative motion ^27^.

### Center of pressure (CoP) testing postural sway

Postural sway was evaluated using the center of pressure (CoP) assay. The mice were weighed and placed on a force plate designed to measure the forces due to postural changes in X, Y, Z, and its angular moments in the XY, YZ, and XZ directions (**Fig 1**). We used the AMTI Biomechanics Force platform (model HEX6x6) and its corresponding AMTI automated acquisition software. An accessory plexiglass cover is placed over the force plate to prevent mice from moving off the force plate. When placed on the force plate, mice are given 2 to 5 minutes to acclimate to the novel environment and minimize their exploratory behavior. After acclimation, ∼10-12 CoP areas were recorded per mouse (resolution per CoP trial = 300 samples per second) and these trials were captured when the mouse showed no active exploratory behavior (e.g., grooming, standing) and its four paws were touching the surface of the force plate. A MATLAB code was used to analyze CoP data, generating a 95% confidence ellipse that enclosed 95% of the CoP trajectory values computed in each of the 1—12 trials. A 10% robust outlier removal (ROUT) was applied to eliminate outlier CoP areas from each individual mouse trials. After this exclusion, each mouse had a minimum of six CoP areas available for averaging. No individual mice were excluded from the study. This measure of postural sway exhibits excellent test-retest reliability ^27^.

### Data analysis and statistics

All statistical analyses were conducted in GraphPad Prism 9.5. Analyses were conducted separately in females and males unless otherwise noted. Multivariate ANOVAs and mixed-effect models were used to analyze Δ tail vasodilations and center of pressure (CoP) ellipse areas, and are further elaborated in the results. Group averages of Δ tail vasodilations and CoP ellipse areas are reported as mean ± SEM, and significance was set at p < 0.05 for all analyses.

Analysis of Δ tail vasodilations involved defining a threshold of 1.5°C to examine the data as a binary outcome test. Tail temperature changes equal to or greater than +1.5°C were designated a Δ tail vasodilation and those less than +1.5°C did not meet the criteria, and were labeled as diminished tail vasodilations. Mice were excluded from further testing if their Δ tail vasodilation after vehicle (saline) testing did not meet the criteria.

Second-order curve fitting was used for fitting head temperatures (B2*X^2^ + B1*X + B0) and R^2^ fit were calculated per curve. Head recovery (mins) was approximated by normalizing head temperature at x = 0 mins to y = 0 °C. An x-intercept quadratic model approximated the recovery by detecting x-intercepts passed 20 minutes (when we stopped the perturbation).

## Results

### No difference in Motion-Induced Δ Heads/Magnitude Hypothermia or Recovery between nestin/hRAMP1 and unaffected controls

Head temperatures are depicted after intraperitoneal (IP) delivery of vehicle (saline), 1.0x CGRP, 1.0x CGRP + 1.0x Olcegepant, and 1.0x CGRP + 1.0x rizatriptan in males and females (**Fig. 2A-J**). The magnitude of hypothermia was computed during an animal’s test – designated as the Δ head – and a 2-way ANOVA was computed across the factors i) treatment and ii) strain in males and in females. No differences in hypothermia magnitude were observed in mice regardless of treatment **(Fig. A-J).** In addition, no differences in recovery were observed in the nestin/hRAMP1 mice compared to their controls for either sex (**Table 1**). This suggests that hypersensitivity to CGRP is not affected by any changes in head temperature drop during or after rotation.

**Fig. 2:**
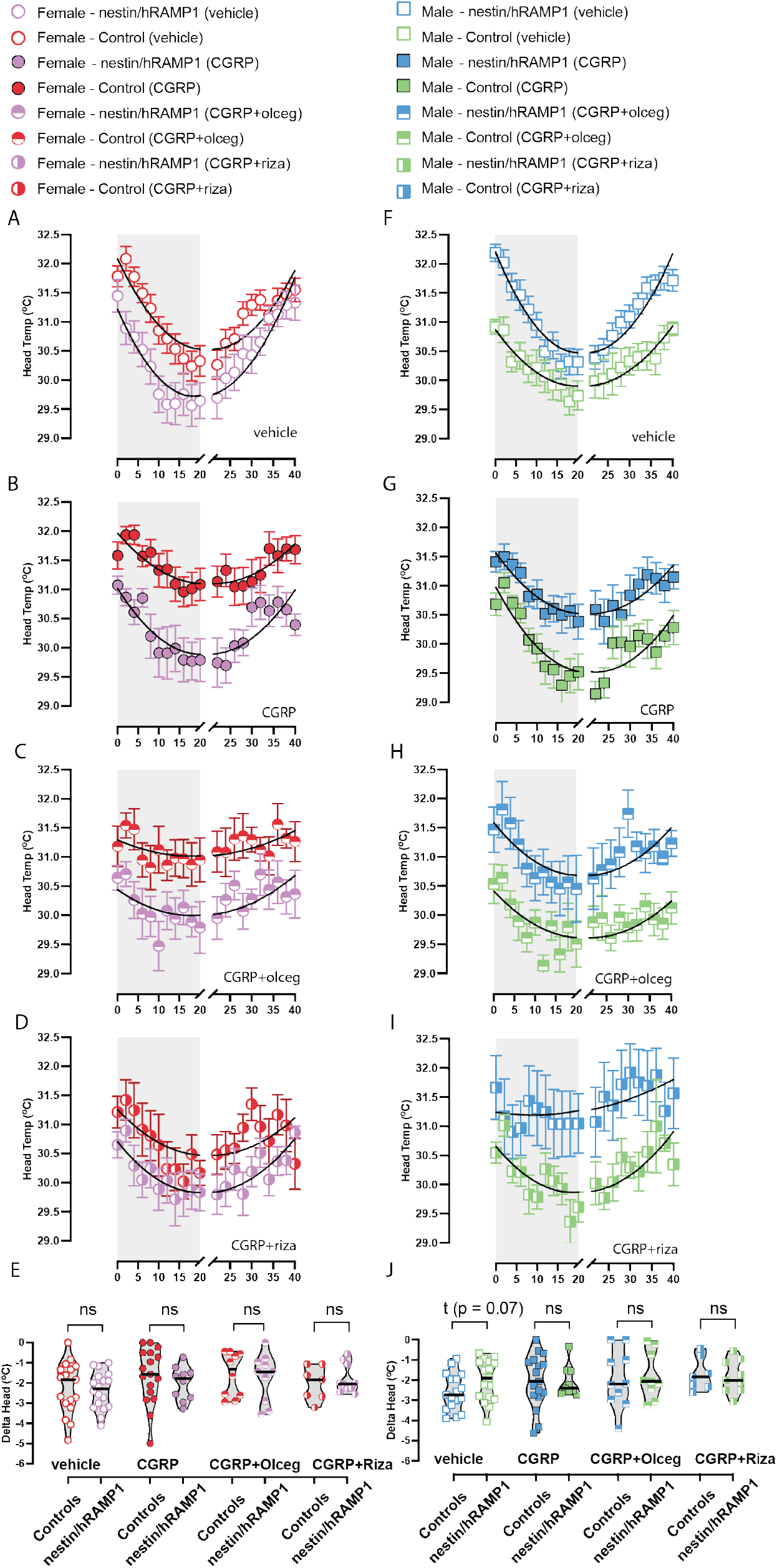
Head temperatures during motion-induced nausea test after intraperitoneal (IP) injections of vehicle, CGRP, CGRP + olcegepant, or CGRP + rizatriptan in female **(A – D)** and male **(F-I)** nestin/hRAMP1 vs controls**. (E-J)** Hypothermia magnitudes (Δ heads) were not different between nestin/hRAMP1 and controls after each treatment in either sex. Sample sizes are listed for male (M) and female (F): vehicle = 17M/18F nestin/hRAMP1 and 27M/26F controls; CGRP - 9M/10F nestin/hRAMP1 and 17M/17F controls; CGRP + Olceg - 9M/10F nestin/hRAMP1 and 10M/10F controls; CGRP + Riza - 9M/10F nestin/hRAMP1 and 7M/7F controls. A legend is provided to indicate color code and symbol shape across treatment, mouse strain, and sex.

**Table 1.** Time to recover from hypothermia (minutes) was analyzed between nestin/hRAMP1 and controls using an x-intercept quadratic model. Males and females were separated. A constraint to the model was applied so that the x-intercept analyzed occurred after time = 20 minutes. While F-statistics and p-values are listed, comparisons are made more confidently in curve fits with R^2^ ≥ 0.60. No significant differences were observed in hypothermia recovery between nestin/hRAMP1 and controls. While 1.0x CGRP + 1.0x olcegepant testing appeared significant between female nestin/hRAMP1 and controls, curve fits were poor so an interpretation is not made. Sample sizes are listed for male (M) and female (F): vehicle = 17M/18F nestin/hRAMP1 and 27M/26F controls; CGRP - 9M/10F nestin/hRAMP1 and 17M/17F controls; CGRP + Olceg - 9M/10F nestin/hRAMP1 and 10M/10F controls; CGRP + Riza - 9M/10F nestin/hRAMP1 and 7M/7F controls/

| Recovery from Hypothermia - controls vs nestin/hRAMP1 |  |  |  |  |  |  |  |  |
| --- | --- | --- | --- | --- | --- | --- | --- | --- |
| Sex | Treatment | nestin/hRAMP1 |  | controls |  | Recovery Difference | F(DFn, DFd) | p-val |
|  |  | Recovery | Fit | Recovery | Fit |  |  |  |
| Male | vehicle | 40.10 | 0.90 | 39.80 | 0.88 | 0.30 | 0.08056 (1, 36) | 0.78 |
|  | 1.0x CGRP | 42.00 | 0.73 | 40.63 | 0.82 | 1.37 | 0.2664 (1, 36) | 0.61 |
|  | 1.0x CGRP + 1.0x Olceg | 44.00 | 0.50 | 39.60 | 0.50 | 4.40 | 1.598 (1, 36) | 0.21 |
|  | 1.0x CGRP + 1.0x Riza | 35.94 | 0.55 | 36.33 | 0.36 | -0.39 | 0.01133 (1, 36) | 0.92 |
| Female | vehicle | 39.30 | 0.79 | 38.28 | 0.90 | 1.02 | 0.7515 (1, 36) | 0.39 |
|  | 1.0x CGRP | 40.70 | 0.67 | 40.80 | 0.83 | -0.10 | 0.001123 (1, 36) | 0.97 |
|  | 1.0x CGRP + 1.0x Olceg | 39.39 | 0.39 | 31.50 | 0.32 | 7.89 | 3.294 (1, 36) | 0.08 |
|  | 1.0x CGRP + 1.0x Riza | 38.79 | 0.74 | 41.44 | 0.34 | -2.65 | 0.7528 (1, 36) | 0.39 |

### CGRP dose response on motion-induced tail vasodilations in nestin/hRAMP1 mice

Motion-induced nausea was examined at three different IP doses of CGRP – 0.01 mg/kg, 0.05 mg/kg, and 0.1 mg/kg. These doses correspond to 0.1x CGRP, 0.5x CGRP, and 1.0x CGRP respectively. When mice are injected with vehicle (saline), mice exhibit a transient delta tail vasodilation in response to the vestibular perturbation. However, this response is diminished after 1.0x CGRP (**Fig. 3B, E**). Dunnett multiple comparison’s test was used to compare Δ tail vasodilations changes after CGRP dose to vehicle, and a significant effect on was observed after intraperitoneal injections of 0.5x CGRP and 1.0x CGRP (**Fig. 3C, F**). Of the male controls, 50% and 53% exhibited diminished tail vasodilations after treatment of 0.5x and 1.0x CGRP (*adj. p = 0.1 and adj. p < 0.001 respectively*), whereas 100% and 83% of male nestin/hRAMP1 mice were affected by 0.5x CGRP and 1.0x CGRP (*adj. p = 0.004* and *adj. p = 0.02 respectively)*, highlighting a potential sensitivity to CGRP in the male nestin/hRAMP1. Most female controls (78% and 88%) exhibited diminished Δ tail vasodilations with doses of 0.5x and 1.0x CGRP (*adj. p < 0.0001 at each dose*), whereas 88% and 80% of female nestin/hRAMP1 tested at 0.5x CGRP and at 1.0x CGRP exhibited diminished Δ tail vasodilations (*adj. p = 0.0007 and adj. p = 0.001 respectively*). Based on the dose response experiments, it is evident that titrating the dose of CGRP to 1.0x leads to disruption in motion-induced nausea in a strong majority of mice. It is also observed that nestin/hRAMP1 males are more readily affected by incremental doses of CGRP than their complement controls (**Table 2**).

**Fig. 3:**
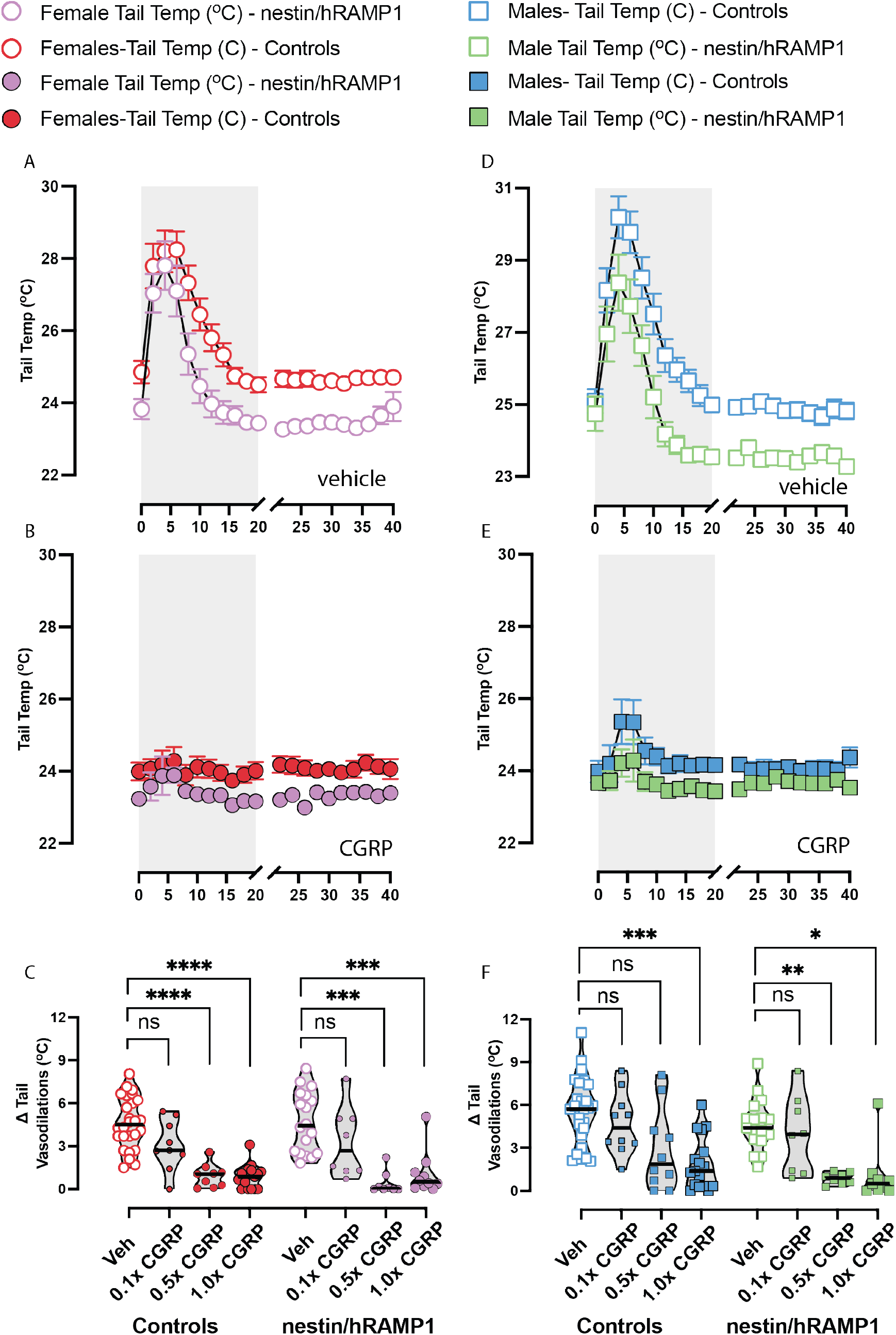
Tail temperatures during motion-induced nausea test and dose response analyses of Δ tail vasodilations are depicted in females (left) and males (right). **(A, D)** Mice exhibit a transient tail vasodilation approximately 5 to 10 minutes after the rotation starts. However, this response is diminished after 1.0x CGRP as seen in **(B, E**). Dunnett multiple comparison’s test is used to compare Δ tail vasodilations at each CGRP dose to vehicle. **(C)** In female nestin/hRAMP1 and controls, a significant effect on Δ tail vasodilations is observed due to 0.5x CGRP and 1.0x CGRP. **(F)** Male controls were affected by 1.0x CGRP but not 0.5x CGRP, whereas male nestin/hRAMP1 mice were affected by 0.05x CGRP and 1.0x CGRP, highlighting a potential sensitivity to CGRP in the male nestin/hRAMP1. Sample sizes are listed for male (M) and female (F): vehicle = 17M/18F nestin/hRAMP1 and 27M/26F controls; 0.1x and 0.5x CGRP = 8M/8F nestin/hRAMP1 and 10M/9F controls, 1.0x CGRP - 9M/10F nestin/hRAMP1 and 17M/17F controls. A legend is provided, and significance levels are indicated as follows: p < 0.05*, p < 0.01**, p < 0.001***.

**Table 2.** Percentage of mice with diminished delta tail vasodilations during motion-induced thermoregulation test after each treatment (row). Columns are categorized by nestin/hRAMP1 vs controls and male vs female.

| ***Diminished $\Delta$ tail vasodilations (< 1.5 ° C)*** |  |  |  |  |
| --- | --- | --- | --- | --- |
| **Treatment** | **Controls (F)** | **nestin/hRAMP1(F)** | **Controls (M)** | **nestin/hRAMP1 (M)** |
| **Vehicle** | 0% | 0% | 0% | 0% |
| **0.1x CGRP** | 22% | 38% | 0% | 38% |
| **0.5x CGRP** | 78% | 88% | 50% | 100% |
| **1.0x CGRP** | 88% | 80% | 53% | 89% |
| **0.5x CGRP + 1.0x Olceg** | 14% | 38% | 20% | 38% |
| **1.0x CGRP + 1.0x Olceg** | 60% | 70% | 20% | 33% |
| **1.0x CGRP + 1.0x Riza** | 71% | 100% | 86% | 89% |
| **0.5x CGRP + 1.0x Riza** | NA | 100% | NA | 100% |

**Table 3.**
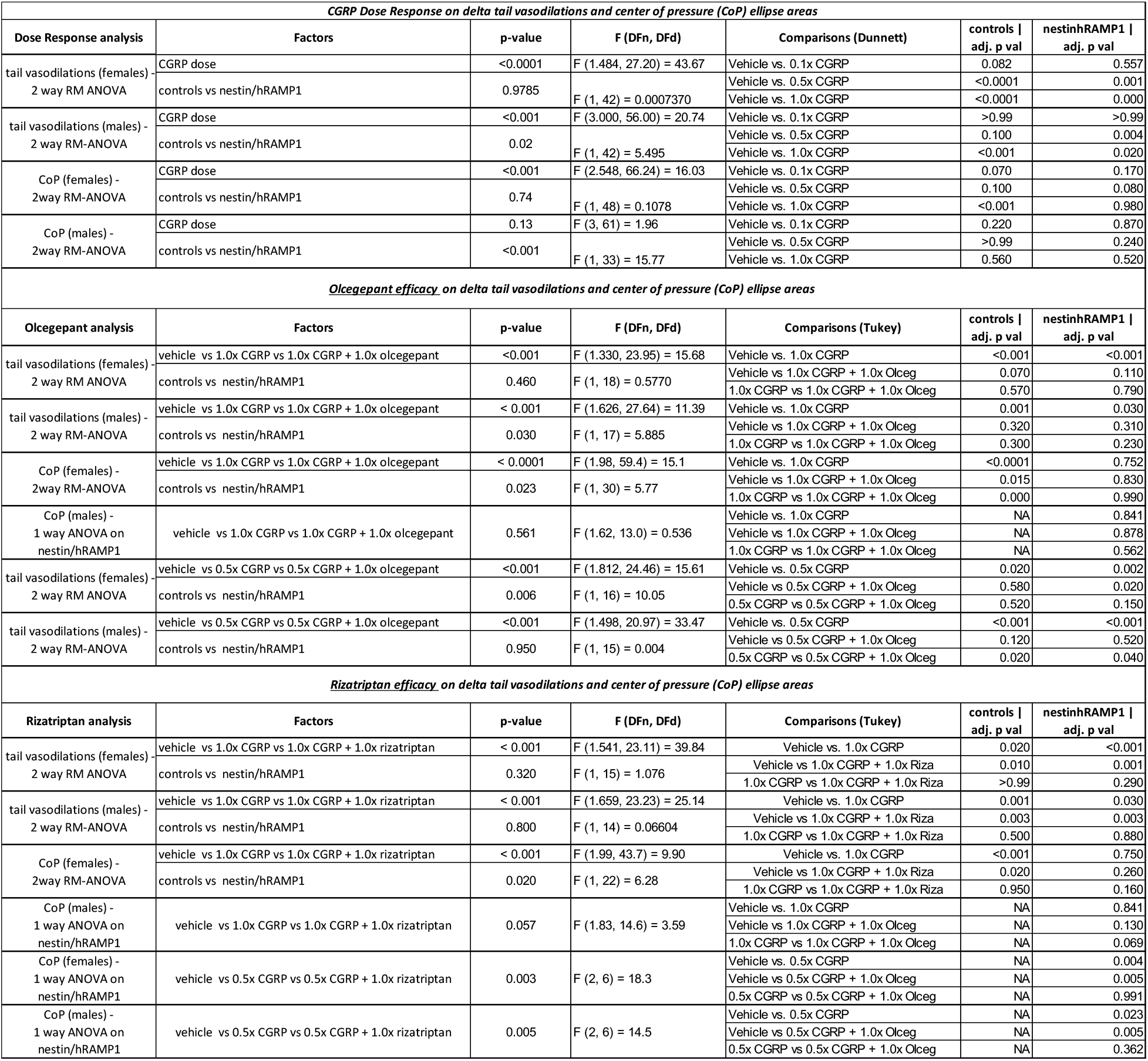
Multivariate statistics are listed per study with corresponding F and p values. Dose responses were analyzed using 2-way repeated measures (RM) mixed effects models to handle missing values since not all animals were repeatedly tested with every CGRP dose, whereas 1.0x CGRP + 1.0x blocker studies used 2-way RM ANOVAs since animals were repeatedly tested.

### Olcegepant’s efficacy on Δ tail vasodilations is similar between nestin/hRAMP1 and controls

The CGRP-receptor antagonist olcegepant was assessed for its efficacy in reducing CGRP’s effect on nausea. Raw temperatures after intraperitoneally delivered (IP) 1.0x CGRP + 1.0x olcegepant treatment are depicted (**Fig. 4A, D**). Post-hoc analysis was conducted using Tukey’s multiple comparisons test. In female controls, 1.0x CGRP caused reduced Δ tail vasodilations, but co-administration of olcegepant helped to block this response in a portion of females. This observation was also seen in nestin/hRAMP1 mice, where some females showed restored Δ tail vasodilations with olcegepant (**Fig. 4B**). In males, olcegepant was effective in blocking 1.0x CGRP-induced changes in male controls and male nestin/hRAMP1 mice (**Fig. 4E**). A separate group of mice were tested at 0.5x CGRP, and mice with diminished Δ tail vasodilations were further tested with 0.5x CGRP + 1.0x olcegepant. We observed a majority of CGRP-affected female controls and nestin/hRAMP1 to regained their delta tail vasodilations with olcegepant blocking this sub- maximal dose. However, unlike male controls where a majority regained their tail vasodilation with olcegepant, male nestin/hRAMP1 still presented with diminished Δ tail vasodilations compared to their vehicle response (*0.5x CGRP vs 0.5x CGRP + 1.0x Olceg, p = 0.04*), suggesting an enhanced sensitivity to CGRP at this submaximal dose that is not being blocked with olcegepant (**Fig. 4F**).

**Fig. 4:**
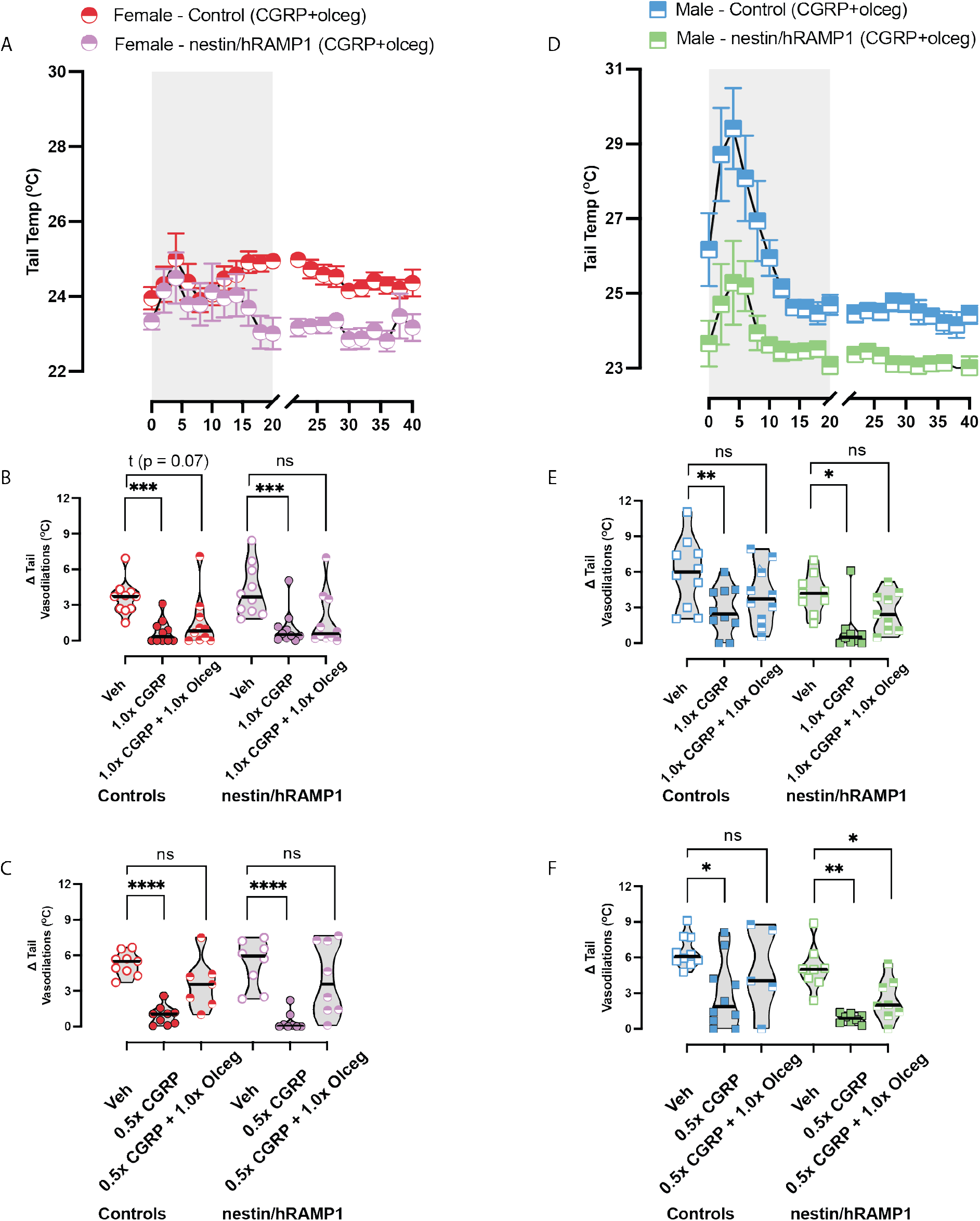
The CGRP-receptor antagonist olcegepant was assessed for its efficacy in motion- induced nausea. Drugs were injected intraperitoneally at the following concentrations: 0.5x CGRP – 0.05 mg/kg, 1.0x CGRP – 0.1 mg/kg, 1.0x olcegepant – 1 mg/kg. **(A, D)** Tail temperatures after intraperitoneally delivered CGRP + olcegepant treatment are depicted. Post-hoc analysis was conducted using Tukey’s multiple comparisons test. **(B)** In female controls, 1.0x CGRP caused reduced Δ tail vasodilations, but co-administration of olcegepant helped some female controls regain a similar response to their vehicle test. This observation was also seen in nestin/hRAMP1 mice, where some females showed restored Δ tail vasodilations with olcegepant. (E) Olcegepant was more effective in blocking CGRP- induced changes in male controls and male nestin/hRAMP1 mice. A separate group of mice were tested at 0.5x CGRP, and mice with Δ tail vasodilations < 1.5 °C were further tested with 0.5x CGRP + 1.0x olcegepant. **(C)** A majority of CGRP-affected female controls and nestin/hRAMP1 regained their tail vasodilation response with olcegepant blocking this sub- maximal dose. **(F)** While a majority of 0.5x CGRP-affected male controls regained their tail vasodilation with olcegepant, many male nestin/hRAMP1 still presented with diminished Δ tail vasodilations compared to their vehicle response. For detailed F-statistics and p-values, please refer to Table 2. Sample sizes are listed: vehicle, 1.0x CGRP, and 1.0x CGRP + 1.0x olcegepant - 9M/10F nestin/hRAMP1 and 10M/10F controls; 0.5x CGRP = 8M/8F nestin/hRAMP1 and 10M/9F controls; 0.5x CGRP + 1.0x olcegepant = 8M/7F nestin/hRAMP1 and 5M/7F controls. A legend is provided, and significance levels are indicated as follows: p < 0.05*, p < 0.01**, p < 0.001***, p < 0.0001****

### Rizatriptan does not block CGRP-induced changes on Δ tail vasodilations

Rizatriptan was also examined across Δ tail vasodilations in a similar testing scheme and analysis as olcegepant. Delta tail vasodilations during CGRP+rizatriptan testing resembled CGRP-induced changes rather than vehicle in all mice tested. Unlike olcegepant, rizatriptan had no protective effect on Δ tail vasodilations in control or nestin/hRAMP1 mice (**Fig. 5A-D**). We also assessed a subset of nestin/hRAMP1 mice after IP delivery of 0.5x CGRP + rizatriptan (**Fig. 5E**). Even at the half dose, rizatriptan did not recover Δ tail vasodilations in these mice. This finding points to rizatriptan’s failure to serve as a therapy for motion sensitivity of migraine and VM.

**Fig. 5:**
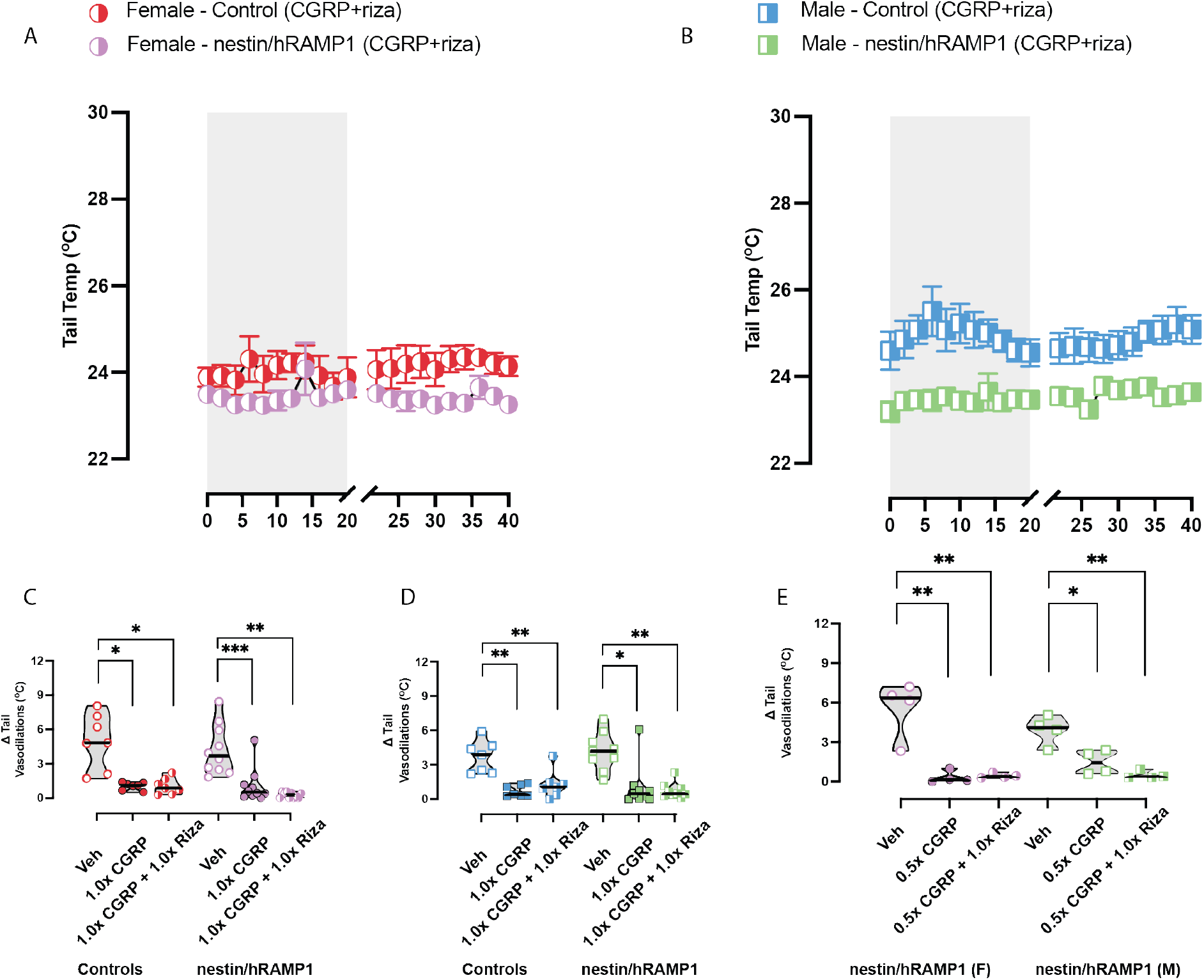
The selective serotonin receptor agonist rizatriptan – a clinically prescribed migraine drug – was assessed for its efficacy in motion-induced nausea. Drug concentrations are listed: 0.5x CGRP – 0.05 mg/kg, 1.0x CGRP – 0.1 mg/kg, 1.0x rizatriptan – 0.6 mg/kg. **(A-B)** Raw temperatures after CGRP + rizatriptan treatment are depicted. **(C - D)** In female and male controls, CGRP caused diminished Δ tail vasodilations and CGRP+rizatriptan did not block these effects. The nestin/hRAMP1 mice resembled control mice, as rizatriptan was not effective in blocking CGRP-induced changes. **(E)** A small subset of nestin/hRAMP1 mice assessed rizatriptan’s efficacy at half the maximum CGRP dose, but no blocker effects were observed. F-statistics and p-values can be found in Table 2. Sample sizes are as follows: vehicle, 1.0x CGRP, and 1.0x CGRP + 1.0x rizatriptan – 9/10 Nestin/hRAMP1 and 7/7 Controls; vehicle, 0.5x CGRP, and 0.5x CGRP + 1.0x rizatriptan – 4F/4M nestin/hRAMP1. Significance levels are listed: p < 0.05*, p < 0.01**, p < 0.001***

### Nestin/hRAMP1 mice exhibit greater postural sway at baseline

Prior to testing CGRP and the effects of blockers, baseline center of pressure (CoP) responses were compared between nestin/hRAMP1 and controls. Two-way ANOVA with Dunnett’s multiple comparisons test computed the following factors: i) male vs female and ii) controls versus nestin/hRAMP1 on sway obtained after vehicle injections (saline). While biological sex had no effect on sway (F (1, 81) = 2.609, p = 0.11), a significant difference was observed between controls and nestin/hRAMP1 (F (1, 81) = 35.45, p < 0.001). Female controls exhibited a CoP of 0.91 ± 0.09 cm^2^, whereas female nestin/hRAMP1 mice had CoPs of 1.56 ± 0.12 (*adj. p < 0.001*). Similarly, male controls had CoP 0.69 ± 0.06 cm^2^, but male nestin/hRAMP1 had larger CoP at 1.41 ± 0.17 (*adj. p < 0.001*). The higher sway in the nestin/hRAMP1 suggest endogenous CGRP having a greater impact on static imbalance due to upregulated hRAMP1 in the nervous system (**Fig. 6A**).

**Fig. 6:**
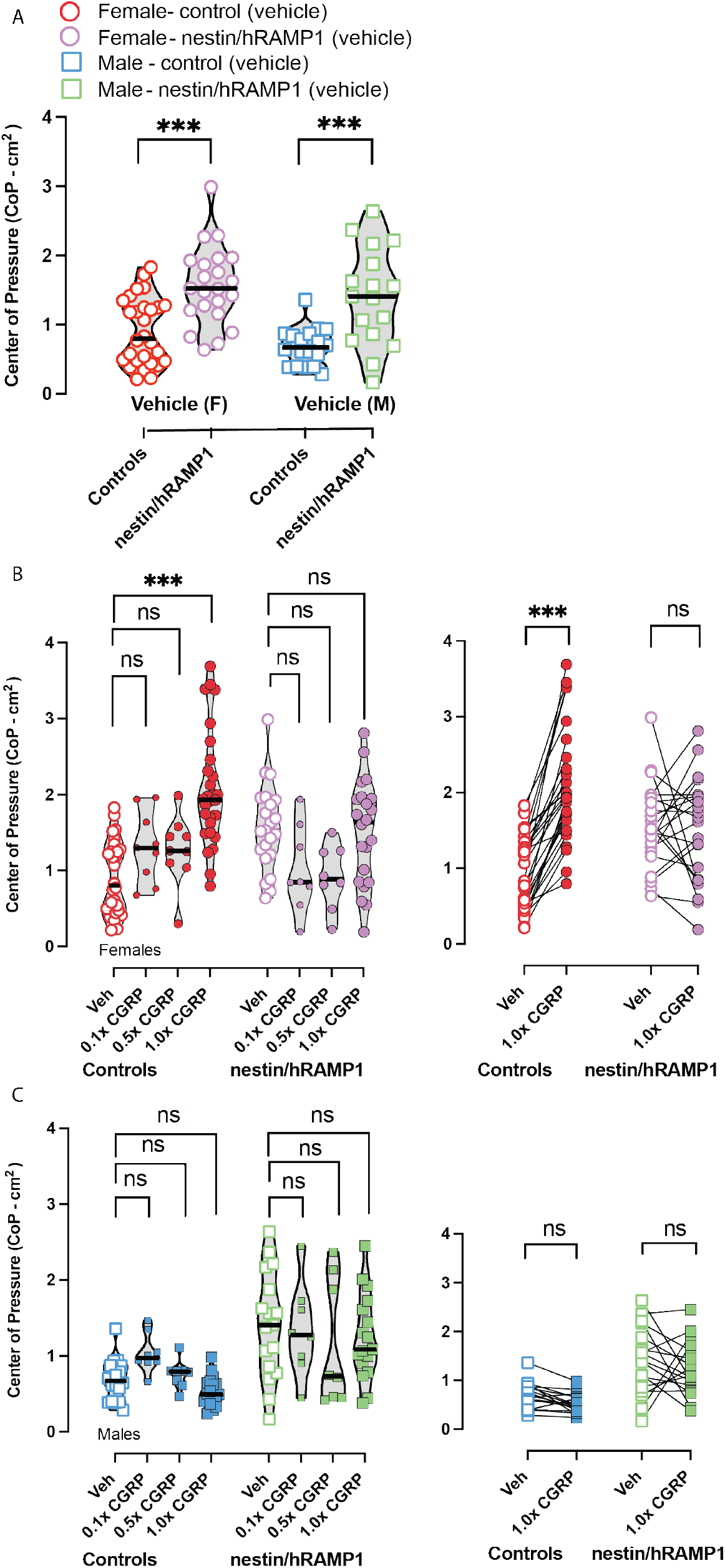
(A) 2-way ANOVA computed the factors i) male vs female and ii) controls versus nestin/hRAMP1 on sway obtained after vehicle injections (saline). While biological sex had no effect on sway (F (1, 81) = 2.609, p = 0.11), a significant difference was observed between controls and nestin/hRAMP1 (F (1, 81) = 35.45, p < 0.001). Tukey’s multiple comparisons indicated nestin/hRAMP1 males and females have a higher sway baseline than their complement controls. **(B)** CGRP induces higher CoP ellipse areas in female controls at 1.0x CGRP but not in female nestin/hRAMP1. **(C)** While male nestin/hRAMP1 sway is significantly higher than male controls at all doses of CGRP, there is no modulation in their sway area with increasing CGRP dose. Before/after plots are placed next to violin plots and show animals that were specifically tested with 1.0x CGRP. Sample sizes are listed: vehicle and 1.0x CGRP– 18M/22F nestin/hRAMP1 and 17M/28F controls; 0.1x and 0.5x CGRP -8M/8F nestin/hRAMP1 and 8M/9F controls. Dose response statistics for sway can be found in top half of Table 2. p < 0.05*, p < 0.01**, p < 0.001***

### CGRP IP injections induce significant sway in female controls but do not increase sway in nestin/hRAMP1 mice

Changes in center of pressure (CoP) after intraperitoneally administered 0.1x CGRP, 0.5x CGRP, and 1.0x CGRP were examined using the same statistical tools used for the motion-induced thermoregulation experiments. Two-way mixed-effected models were used to assess CGRP dose and controls versus nestin/hRAMP1, and Dunnett’s multiple comparisons was used to compare differences in CoP after each CGRP dose to vehicle. In male nestin/hRAMP1 and controls, no differences were detected due to CGRP (**Fig. 6C**).

Female controls exhibited significantly higher CoP after 1.0x CGRP compared to their vehicle response (*adj. p < 0.001*) (**Fig. 6B**). In contrast, female nestin/hRAMP1 mice did not significantly change their CoP in response to increasing doses of CGRP. This suggests that endogenous CGRP in these hypersensitized mice was saturated.

### Olcegepant and not rizatriptan blocks CGRP-induced sway changes in female control mice, but does not block endogenous sway in nestin/hRAMP1 mice

Olcegepant and rizatriptan were examined for their ability to protect against 1.0x CGRP-induced changes in CoP or to protect from endogenous CGRP release in nestin/hRAMP1 mice. Olcegepant led to lower CoP compared to 1.0x CGRP alone (*female controls: CGRP vs CGRP+olcegepant, adj. p = 0.0004*) (**Fig. 7A**). In contrast, rizatriptan did not have an effect on female control CoP, as 1.0x CGRP + 1.0x rizatriptan led to sway that resembled 1.0x CGRP induced changes (**Fig. 7C**). Interestingly, the CoP results on female controls dramatically differ from their nestin/hRAMP1 counterparts. Female nestin/hRAMP1 mice did not further increase their sway after 1.0x CGRP, nor did co-or separate delivery of 1.0x olcegepant or 1.0x rizatriptan modulate their CoP any further.

**Fig. 7:**
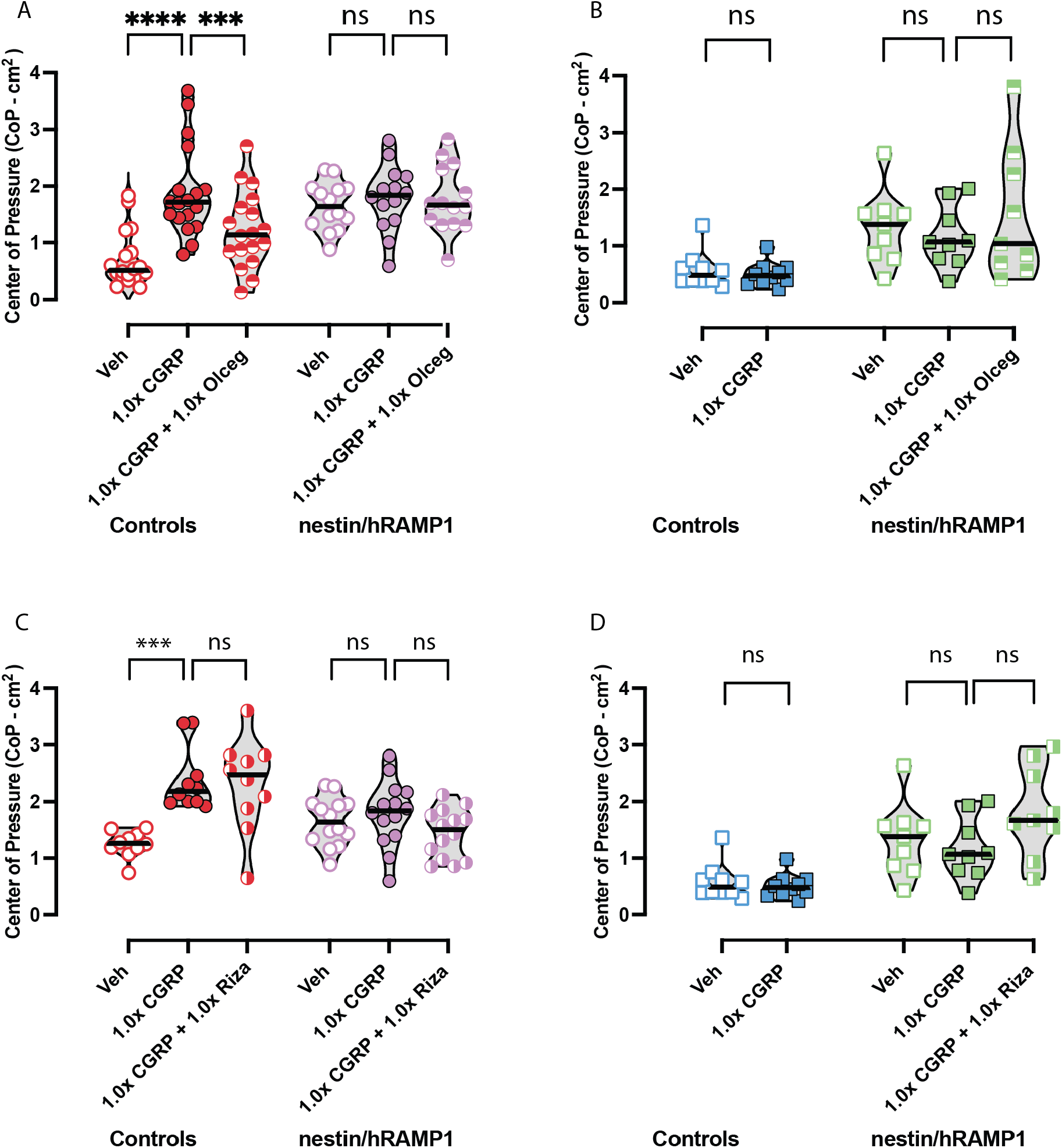
Changes in sway were observed after vehicle, CGRP, CGRP + olcegepant or CGRP + rizatriptan treatment in controls and nestin/hRAMP1, using same concentrations from motion-induced thermoregulation studies. **(A)** In female controls, CGRP causes increases in postural sway but co-administration of olcegepant blocks CGRP-induced changes. In contrast, female nestin/hRAMP1 mice do not appear affected by CGRP, and olcegepant is not an effective blocker. **(B)** Male nestin/hRAMP1 mice do not respond to CGRP or CGRP+olcegepant. For male controls, a paired t-test was used to compare vehicle vs CGRP effects but no differences were observed. The lack of a CGRP-induced change in male control sway provided us the rationale to discontinue testing these mice with blockers. Sample sizes include 9M/14F nestin/hRAMP1 and 10M/18F controls repeatedly tested. Full list of F and p-values for these sway experiments are found in Table 2. Significance levels are indicated as p < 0.05*, p < 0.01**, p < 0.001***, p < 0.0001****.

When examining male controls and nestinhRAMP1 (**Fig. 7B and 7D**), CGRP and CGRP co-delivered with olcegepant/rizatriptan did not modulate their sway when compared to their response to vehicle. Male nestin/hRAMP1 mice have higher CoP values after vehicle and 1.0x CGRP than male controls, but olcegepant and rizatriptan do not affect male CoP sway. **Table 2** shows specific details regarding the multivariate statistics (F-statistics, p- values, and post-hoc analyses) used in this study for motion-induced nausea and postural sway comparisons.

## Discussion

Migraine treatment aims not only to address pain and ensure freedom from headache, but also to alleviate the most bothersome symptoms (MBS), such as nausea, vertigo, photophobia, or phonophobia. Photophobia is the most prevalent MBS and affects 50% of migraine patients, while nausea -the second most frequent MBS- affects 30% of migraine patients ^29^. Furthermore, individuals suffering from migraines also encounter difficulty with postural control and balance, resulting in increased sway area and velocity ^3, 4, 30, 31^.

A study examining various lighting conditions during bipedal and unipedal stances of migraine patients concluded that migraine-associated photophobia is linked to postural sway^32^. Additionally, nausea is linked to calcitonin gene-related peptide (CGRP; and sensory pathways linked to nausea involve the vestibular system and the area postrema, which is a circumventricular organ associated with nausea and vomiting ^33^. CGRP receptors have been detected in the area postrema through the localization of RAMP1 mRNA in this region^34^. As the area postrema is not enclosed by the blood-brain barrier, systemic delivery of CGRP can affect area postrema neurons and potentially alter this region during motion- induced nausea. Moreover, overexpression of RAMP1 may exacerbate this condition.

The nestin/hRAMP1 mouse model has been extensively studied to manipulate CGRP’s effects, simulate migraine-like surrogate behaviors, and assess the efficacy of migraine drugs. The model first elucidated that the CGRP receptor is a critical aspect of neurogenic inflammation ^15^. Moreover, systemic IP injection of CGRP has been extensively studied in preclinical rodent models of migraine where it is believed to affect the PNS. For instance, p0eripheral IP injection of CGRP resulted in light-aversive behaviors and can produce spontaneous pain in mice ^35, 36^. In fact, IP CGRP’s effect on light aversion was independent of any vasodilatory effects as CGRP-induced light-aversion was observed even with normalized blood pressure ^37^. This study examined the impact of systemic CGRP on motion-induced nausea and postural sway using the motion-induced thermoregulation nausea surrogate and center of pressure assays. Systemic IP administration of small- molecule blockers of the CGRP receptor “gepants” have been shown to enter the PNS like CGRP and triptans ^38^. Because of this we also investigated whether the CGRP-receptor antagonist olcegepant or the selective serotonin receptor agonist rizatriptan could mitigate such changes in the nestin/hRAMP1 and unaffected littermate control mice.

The study revealed several fascinating findings. Initially, the investigation did not identify significant differences in head hypothermia magnitude or time to recover from head hypothermia in mice irrespective of strain or treatment. As a result, it is reasonable to conclude that CGRP and its blockers do not partially shape this element of the nausea response in mouse models. However, IP CGRP had a significant impact on the presentation of Δ tail vasodilations in both male and female mice. At higher doses (0.5x and 1.0x CGRP), the majority of mice, including both nestin/hRAMP1 and control mice, showed reduced Δ tail vasodilations. At the highest dose of 1.0x CGRP, which is 0.1 mg/kg, most mice do not show a significant increase in Δ tail vasodilation after being stimulated with provocative rotation. The strong effect observed at the maximum dose of CGRP prompted us to investigate the impact of the CGRP-receptor antagonist olcegepant and the serotonin receptor agonist (5HT1B/1D) rizatriptan.

Olcegepant effectively restored tail vasodilations in male mice and, to a lesser extent, in female mice, in both nestin/hRAMP1 mice and controls. However, rizatriptan did not offer protection and resulted in changes comparable to those resulting from CGRP- induced alterations. Rizatriptan and other triptans have been extensively researched as drugs that indirectly inhibit the release of CGRP in migraine treatment. Further investigation into the effects of half the maximum dose was conducted through the co-administration of olcegepant or rizatriptan. It was discovered that at half the maximum dose of CGRP, olcegepant improved the response of both nestin/hRAMP1 and control mice; however, it was more effective in control male mice than nestin/hRAMP1 male mice. Rizatriptan did not restore Δ tail vasodilations in nestin/hRAMP1 mice, even at a half-dose of CGRP. It is interesting to note that while rizatriptan was ineffective in motion-induced nausea and sway assays, rizatriptan was effective in reducing light sensitivity and allodynia in control and nestin/hRAMP1 mice ^14, 17, 20^. This points to need to assess motion-sensitivity in addition to other sensory sensitivities of migraine in mouse models.

During the investigation of postural sway, we observed that nestin/hRAMP1 mice showed higher baseline sway levels than control mice. Our previous research has demonstrated that males do not encounter variations in their center of pressure while being exposed to provocation rotation or CGRP. In contrast, female controls treated with 0.1 mg/kg CGRP encounter postural sway. In this study, we found that male nestin/hRAMP1 mice have a higher range of sway than their unaffected littermate controls. This change is presumably associated with increased responsiveness to endogenous CGRP, which influence sway due to elevated signaling mediated by hRAMP1 overexpression. This heightened sensitivity is internally consistent with other behavioral studies in nestin/hRAMP1 mice, which also exhibit light sensitivity to normal room lighting that is not observed in unaffected controls^14, 17, 20^. However, unlike light and touch sensitivity, increased postural sway in the nestin/hRAMP1 was not decreased by either olcegepant or rizatriptan blockers. Moreover, the nestin/hRAMP1 mouse had increased CGRP receptor signaling at birth suggesting that perhaps postural pathways may have been altered during development.

Future studies are planned to determine if a conditional increase in hRAMP1 expression would show similar responses to the CGRP receptor blocker, olcegepant.

In conclusion, mice that are sensitized to CGRP (nestin/hRAMP1) mice show increased endogenous postural sway and are less sensitive to CGRP-receptor blockade in a nausea assay. This nestin/hRAMP1 mouse model has been used to model the light and touch sensitivities of migraine and these motion-sensitivity findings allow us to understand CGRP’s role in motion-sensitivity which may serve to develop new therapies aimed at alleviating these debilitating symptoms in migraine patients.

## Acknowledgments

We would like to thank Abigail Dweh and Laura Nafis for their contributions to the data analysis. We would also like to thank Dr. Andrew Russo for providing the hRAMP1-loxP mice for these studies. All figures were created in Biorender.

